# Revisiting the recombinant history of HIV-1 group M with dynamic network community detection

**DOI:** 10.1101/2021.04.01.438061

**Authors:** Abayomi S Olabode, Garway T Ng, Kaitlyn E Wade, Mikhail Salnikov, David W Dick, Art FY Poon

## Abstract

A new abundance of full-length HIV-1 genome sequences provides an opportunity to revisit the standard model of HIV-1/M diversity that clusters genomes into largely non-recombinant subtypes, which is not consistent with recent evidence of deep recombinant histories for SIV and other HIV-1 groups. Here we develop an unsupervised non-parametric clustering approach, which does not rely on predefined non-recombinant genomes, by adapting a community detection method developed for dynamic social network analysis. We show that this method (DSBM) attains a significantly lower mean error rate in detecting recombinant breakpoints in simulated data (quasibinomial GLM, *P* < 8 × 10^−8^), compared to other reference-free recombination detection programs (GARD, RDP4 and RDP5). Applied to a representative sample of *n* = 525 actual HIV-1 genomes, we determined *k* = 25 as the optimal number of DSBM clusters, and used change point detection to estimate that at least 95% of these genomes are recombinant. Further, we identified both known and novel recombination hotspots in the HIV-1 genome, and evidence of inter-subtype recombination in HIV-1 subtype reference genomes. We propose that clusters generated by DSBM can provide an informative new framework for HIV-1 classification.

## Introduction

Understanding the global epidemiology of human immunodeficiency virus type 1 (HIV-1) is contingent on our ability to accurately reconstruct the origin of the virus [1]. The origin of different lineages of HIV-1 has been traced back to multiple zoonotic transmissions of simian immunodeficiency virus from chimpanzees (SIVcpz) directly or through an intermediate host to human populations in West and Central Africa [2, 3]. SIVcpz, on the other-hand, originated through multiple cross-species transmissions of SIV from other primates to chimpanzees, with extensive recombination between two or more ancestral variants [4]. SIVs in other non-human primate species also have evidence of recombinant origins [5], and at least 13 recombinant breakpoints and 14 host-switch events have been identified among SIV lineages overall [6]. Furthermore, there is phylogenetic evidence that the ancestor of HIV-1 group N was a mosaic combination of the HIV-1/M ancestor and an SIV lineage [2]. Taken together, these findings imply that recombination has been an important evolutionary force in the deep evolutionary history of primate lentiviruses.

Currently, HIV-1 is classified into four major groups (M, N, O, P). Group M, which is responsible for the global pandemic, is further subdivided into nine ‘pure’ subtypes (A-D, F-H, J and K) that are defined by substantial genetic divergence (about 10% to 25%) and/or bootstrap support for monophyletic clades [7]. The first evidence of the existence of recombinant HIV-1 genomes was documented in the late 1980s [8]. HIV-1 exhibits a high rate of recombination that is driven by obligate template switching during the reverse transcription stage of its replication cycle [9] and by co-infection at the cellular [10] and host levels [11]. Consequently, there are presently close to 100 circulating recombinant forms (CRFs) documented by the Los Alamos National Laboratory (http://www.hiv.lanl.gov/), where each CRF comprises a mosaic of two or more pure HIV-1 subtypes that has been sampled from at least three epidemiologically-unlinked individuals [12]. There is also a large number of unique recombinant forms (URFs) that do not meet the latter criterion [13]. Nevertheless, the prevailing view of HIV-1 diversity is that the majority of virus genomes can be classified into one of the subtypes or a relatively small number of CRFs that have reached a high level of global or regional prevalence, such as CRF01 AE (southeast Asia [14]) or CRF07 BC (China [15]). Thus, inter-subtype recombination is considered to be an infrequent event, such that new infections can be routinely classified into one of the subtypes or CRFs by sequencing a specific region of the genome, such as the HIV-1 *pol* gene that encodes the major targets of antiretroviral therapy.

Increasing deployment of whole-genome sequencing technologies for HIV-1 around the world is providing a clearer picture of the prevalence and diversity of recombinant HIV-1 genomes [16]. For instance, molecular clock estimates of the origin of the HIV-1 pandemic [17], or the time to the most recent ancestor (tMRCA), may be inaccurate due to a deep recombinant history, such that a single phylogenetic tree is not an adequate representation of discordant evolutionary histories relating different regions of the virus genome. In previous work [18], we used a sliding-window molecular clock analysis of near full-length HIV-1 genomes to show that different regions of the virus genome yield significantly different estimates of the tMRCA. This result supports the hypothesis that HIV-1 group M (HIV-1/M) may have a deep recombinant origin that involved at least two genome fragments with different evolutionary histories. For example, this result could be explained by the introgression of a genome fragment from an unsampled lineage with a more distant common ancestor to the diversity of the present day [19].

This phylogenetic evidence, together with other findings on the deep recombinant history of SIV [6], supports the hypothesis that the evolutionary history of HIV-1/M can be improved if the conventional framework of established ‘pure’ reference subtypes with limited subsequent recombination is revisited. One of the obstacles to evaluating this hypothesis is that most tools for classifying HIV-1 genomes rely on a reference set of predefined subtypes, such as COMET [20] or SCUEAL [21]. In other words, these are supervised classification methods. We propose that the best way to test whether the early evolutionary history of HIV-1/M is reticulate is to apply an unsupervised clustering method to the global diversity of virus genomes. For example, GARD (genetic algorithm for recombination detection [22]) is a divisive unsupervised method that attempts to find the optimal partition of the sequence alignment that maximizes the joint likelihood. Fitting GARD to large numbers of near full-length genomes is computationally intensive because phylogenies must be reconstructed for each genome segment defined by every candidate partition. Furthermore, a post-processing step is required to determine support for a recombination breakpoint from the topological discordance of phylogenies. In addition, there are a number of recombination detection heuristic methods that do not require reference sequences, several of which are implemented in the software package RDP (recombination detection program) [23]. However, this package is only distributed as binaries compiled for the Windows operating system.

This study endeavours to reconstruct the extent of recombination across the entire HIV-1/M genome. Here we describe a new unsupervised non-parametric approach to this problem based on adapting a community detection method that was developed for the analysis of dynamic social networks [24]. Community detection consists of grouping or partitioning nodes (vertices) of a network graph into the same community based on their relative edge density [25, 26]. We note that a connected component in the network, which is often referred to as a ‘cluster’ in the context of genetic epidemiology [27], may comprise multiple communities. Likewise, network communities are often referred to as clusters. To avoid confusion, we will explicitly refer to subgraphs in which there are no edges to external nodes as connected components instead of clusters.

A network in which the distribution of edges among nodes remains constant over the observational time period is a static network [28]. However, real networks are dynamic as relationships evolve over time, represented by the addition and/or removal of network edges [29, 30]. A dynamic network can be represented by a series of static networks that each capture the state of the system at a given point in time. In the context of a dynamic network, communities are a group of nodes that are stably connected by a relatively high density of edges over time [31]. The development of methods to detect network communities is an active area of research with a broad domain of application, including biological sciences, for extracting patterns from complex relational data [26].

In this study, we use genetic distances between sequences to generate a network or graph, where each node represents a virus genome, and each edge indicates that the respective genomes have a distance below some threshold. We assume that communities in the resulting graph correspond to phylogenetic clusters such as subtypes. When distances are calculated from different regions of the genome, a node may switch membership from one community to another, which can correspond to a recombination event. This is analogous to an individual in a social network switching affiliations from one community to another. We use stochastic block modelling to detect the community structure of the genetic similarity graph, and employ a recently described [32] expectation maximization algorithm to estimate the ‘migration’ rates between communities. Stochastic block models (SBMs) are one of the most widely utilized class of models for community detection in networks [24]. Individuals belong to one of *K* latent communities, and the probability of an edge between individuals is determined only by community membership, *e*.*g*., *P*_*ii*_ *> P*_*i j*_ for *i, j* ∈ {1,…, *K*}. SBMs are designed to uncover hidden structural features in complex networks by clustering nodes based on similar or shared attributes [33], while dynamic stochastic block models (DSBMs) uncover these hidden data structures as a dynamic network changes over discrete time [34]. We propose to adapt DSBMs as an unsupervised method to characterize the effect of recombination on the evolutionary history of HIV-1 genomes.

## Methods

### Data processing

A total of *n* = 3, 900 near full length (>8,000 nt) HIV-1/M genomes, manually curated from our previous study [18] were used in this study. Briefly, we constructed a reduced multiple sequence alignment based on the pairwise alignment of sequences against a consensus genome sequence, discarding insertions relative to this reference to filter out regions of relatively low evolutionary homology. To minimize the computing time of subsequent steps in our analysis, we selected *n* = 550 sequences from this alignment by progressively removing genomes with the shortest genetic distances to other genomes in the dataset. Next, we realigned the remaining sequences using MAFFT (version 7.271) [35], then partitioned the alignment into sliding windows of 500nt at steps of 100nt, resulting to a total of 82 alignment subsets covering HXB2 nucleotide coordinates 790 to 9465. (We arrived at these parameter settings after some preliminary tests varying window and step sizes.) Sequences that had deletions spanning more than 20% in one or more subsets were excluded, resulting in a final total of 525 sequence subsets. We used the SCUEAL (Subtype Classification Using Evolutionary ALgorithms) method implemented in HyPhy to classify these genomes with respect to the HIV-1/M subtype reference sequences obtained from the Los Alamos National Laboratory (LANL) HIV Sequence Database (http://www.hiv.lanl.gov).

### Community detection

The aim of this study was to characterize the extent of recombination over a series of networks as snapshots of the dynamic system over time, throughout the evolutionary history of HIV-1/M. We are adapting DSBMs to this problem by drawing an analogy between time and the length of the HIV-1 genome, with recombination breakpoints as discrete events. If recombination is relatively infrequent, then genomes should largely cluster into pure subtypes that diverge over time from a common ancestor. With abundant recombination, however, we would expect sequences to switch frequently between communities. To construct a graph, we used the *dist*.*dna* function of the R package *ape* [36] to compute the Tamura-Nei [37] (TN93) distances between every pair of sequences within each of the 82 windows extracted from the alignment. We imported the resulting distance matrices into R to generate undirected graphs using the package *igraph* [38].

We decided not to use a fixed threshold distance for all windows because rates of molecular evolution vary substantially from one part of the HIV-1 genome to another. Instead, we applied different quantile thresholds (*e*.*g*., the upper 25%) to the observed distributions of TN93 distances for each window, and used simulation experiments (see below) to assess which quantile yielded the most informative graphs. Finally, to reconstruct the distribution of recombination events in the full alignment, we used the R package *dynsbm* [39] to fit a DSBM to the series of graphs. We used the integrated completed likelihood (ICL) criterion [40] to determine the optimal number of latent groups (*K*). The ICL criterion is often used in model-based clustering applications to select the appropriate number of communities [41, 42].

To predict recombination breakpoints from the DSBM outputs, we used the R package *changepoint* to perform change point detection on cluster assignments along the sequence of windows for each genome. We dropped missing values from each sequence up to a maximum tolerance of 10 missing values. Furthermore, we evaluated both the default ‘at most one change’ (AMOC) method [43] and the more recently described pruned exact linear time (PELT) algorithm [44] with a minimum segment length of 10 windows. In both cases, we used the default modified Bayesian information criterion (MBIC) to penalize the addition of change points [45].

### Recombination simulation

Simulating sequence evolution is an important tool for validating new methods in phylogenetic analysis by evaluating their accuracy on data with known parameters. To calibrate our DSBM method, we evaluated its prediction accuracy on simulations under varying parameter settings. In addition, we compared its performance to other recombination detection methods that do not require reference genomes, *i*.*e*., GARD and RDP. First, we created a basic and simplified recombination scenario by selecting four reference sequence each of ‘pure’ HIV-1/M subtypes curated by LANL http://www.hiv.lanl.gov/ A, B, C, and D (n=16) and aligning them. For each breakpoint category a uniform breakpoint location and fragment was selected and concatenated for both parent and child/children (recombinant) trees as follows: 4500nt, 3000nt, 2250nt for 1, 2 and 3 breakpoints respectively. We then evaluated our ability to reconstruct these simulated recombination breakpoints. This ‘post-hoc’ simulation method is similar to the one used to validate the subtyping and recombination detection program COMET [20], which is reference-dependent. Because one of our objectives was to screen the full diversity of HIV-1 for recombination, we repeated this first simulation method for a larger set (*n* = 37) of subtype reference sequences including representatives of subtypes A-H, J and K.

Furthermore, we generated more realistic simulations in which recombination events could be distributed throughout the evolutionary history of the ‘observed’ sequences. First, we used IQ-TREE (version 1.3.11.1) [46] to reconstruct a maximum likelihood tree relating the subtype reference sequences used in the previous method. We rooted this tree using the sample collection dates by root-to-tip regression using the *rtt* function in the R package *ape* [36]. Next, we used BEAST (version 1.10.4) [47] to sample time-scaled phylogenies from the posterior distribution, using the Tamura-Nei (TN93) nucleotide substitution model, with rate distribution modeled by a gamma distribution with 4 rate categories; an uncorrelated lognormal clock model (*µ* = 1, *σ* = 0.33); and a skyline tree prior with 10 population sizes. In addition, we set the prior distribution for the time to the most recent common ancestor to a normal distribution with *µ* = 82 years and *σ* = 4.1, based on recent estimates of the origin of HIV-1/M [1]. We ran a single chain sample for 10^8^ steps, discarded the first 10^7^ steps as burn-in, and generated a maximum clade credibility (MCC) tree from the remainder using TreeAnnotator (version 1.10.4).

We used a Python script to update the MCC tree with recombination events by switching random nodes that span a randomly selected time. To generate 1, 2 or 3 recombination breakpoints, we selected a ran-dom point in time to prune and regraft subtrees on two randomly selected extant branches. We assumed that the leftmost portion of the alignment was always related by the original tree, and that the portion of the alignment past the first breakpoint was related by the modified tree. We repeated this process for additional breakpoints by progressively modifying the tree to the immediate left of the breakpoint to relate sequence fragments to the right. In all, we generated 100 simulations of 1, 2 and 3 breakpoints for a total of 300 trees. The evolution of a 9,000 nt sequence at the root was simulated along each tree using INDELible (v1.0.3) [48] under a codon substitution model with rate variation modeled by a gamma distribution (*α* = 1.5, *β* = 3) discretized into 50 rate categories, and a transition/transversion ratio *κ* = 8.0. The input tree was rescaled such that the expected number of substitution events per codon was 3.22, which we derived from the total length of the maximum clade credibility tree (643.5 years), clock rate estimate (0.00167 substitutions/nt/year) and an adjustment of 3 nucleotides per codon.

We used the sequence alignments produced by both simulation methods to evaluate different recombination detection methods, including the DSBM community detection method. We ran GARD (genetic algorithms for recombanation detection) method in HyPhy (version 2.3.11) [22] with the HKY85 nucleotide substitution model and no rate variation. Since this program required a message passing interface (MPI) parallel computing environment, we ran this analysis with 8 threads. In addition, we evaluated both versions 4 and 5 of the program RDP (recombination detection program) [23, 49] using the default settings and with the sequence type set to ‘linear’.

To measure the computing times of these different methods, we generated test sets by randomly sampling sequences with replacement from the larger set of 37 reference genomes, and added random mutations at 0.1% of positions in each genome on average. Through this process, we generated 10 replicates of 50, 100 and 200 sequences for a total of 30 test sets, and then ran each set through DSBM, GARD, RDP4 and RDP5. Since the RDP programs are only released as Windows binary executables, and GARD requires an MPI-enabled environment, it was not feasible to run all tests in the same computing environment. Hence, we focused on characterizing the time complexity of each method (relative change in time with increasing data).

## Results

Our hypothesis is that community detection with a dynamic stochastic block model (DSBM) [32] can be adapted for the unsupervised detection of recombination events from variation among HIV-1 genomes. We draw an analogy between latent communities in a social network and genetic clusters, such as the clusters that are presently labeled as the HIV-1 subtypes. Hence, a recombination event is analogous to moving from one network community to another. To briefly summarize our approach, we first generate a series of networks by partitioning a multiple alignment of HIV-1 genomes into ‘sliding windows’ of a fixed width and step size, and then compute a genetic distance (TN93) between all pairs of sub-sequences within each window. A network (undirected graph) can be derived from each distance matrix by applying a distance threshold, below which an edge is drawn between the respective nodes (sequences). Finally, we applied the implementation of DSBMs in R [39] to the resulting series of graphs to parameterize the transition matrix modelling the movement of individuals between network communities. For brevity, we will refer to this method as DSBM.

### Simulation analysis

To evaluate the accuracy of this approach, we first used a simple method to generate alignments of recombinant sequences by combining fragments from one or more HIV-1/M subtype reference genomes at pre-determined positions (breakpoints). We also used this simulation experiment to optimize the threshold for converting TN93 distances to graphs. DSBM performed best at a TN93 threshold of 40%, where the recombinant fragments were correctly identified in 12 out of 15 replicate simulations with an average accuracy of 99% (Supplementary Figure S1). The remaining 3 of the 15 simulations consist of recombination between subtypes B and D. DSBM did not detect recombination events between subtypes B and D, simply because the method tends to classify subtypes B and D sequences as one group. Next, we generated more realistic simulations by grafting recombination events into a time-scaled phylogeny reconstructed from HIV-1 genomes with prior information [1]. Starting from the left of the alignment, we pruned and regrafted subtrees by selecting two extant branches at a random at given time points to produce a new tree for sequence fragments past the next breakpoint. We tested the ability of DSBM to accurately capture the recombination breakpoints in these simulations (Figure 1A). Overall, DSBM was significantly more accurate (measured by error percentage) than the three other methods on these data (quasibinomial GLM, *t >* 5.4, *P* < 7.5 *×* 10^− 8^). Error rates increased significantly with the number of actual recombination breakpoints in the data (*t* = 15.6, *P* < 10^− 12^).

**Figure 1:**
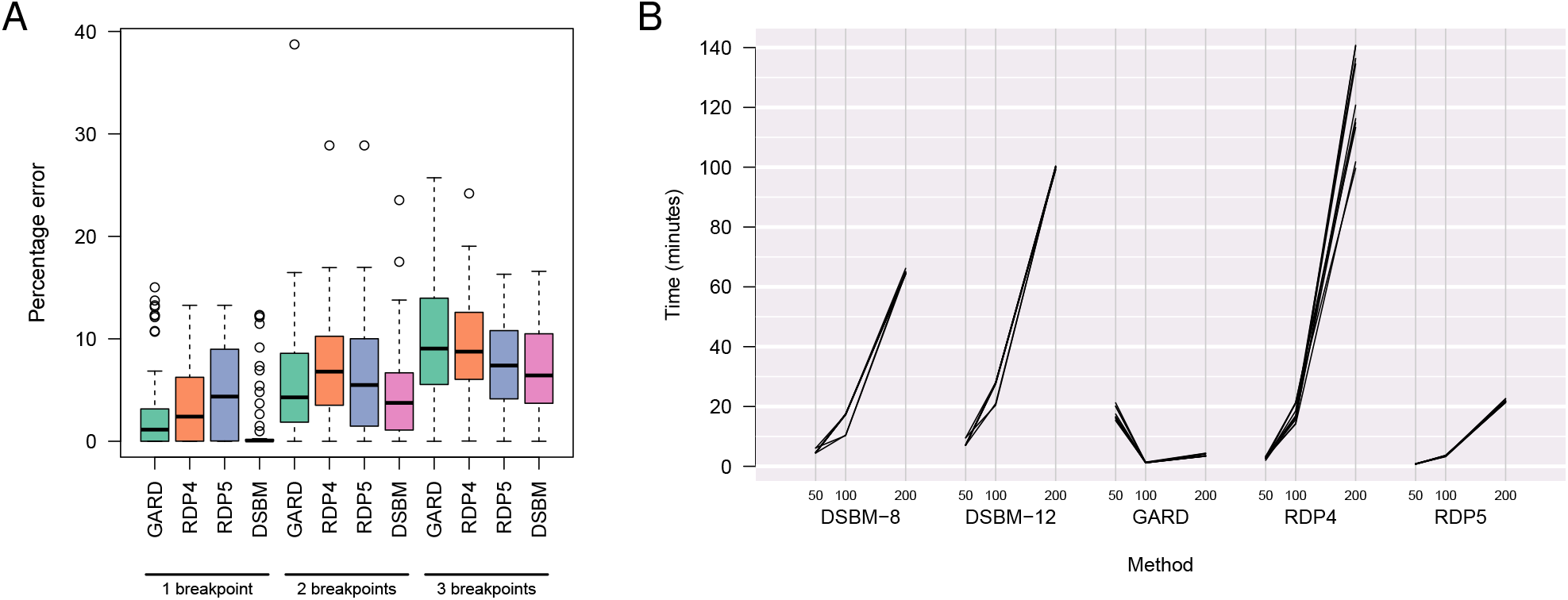
(A) Box-and-whisker plots summarizing the percent error of recombination detection methods on sequences simulated from a time-scaled HIV-1/M phylogeny. Percent error was calculated by mapping outputs from each program to the same outcome space of cluster assignments by sequence and nucleotide position. (B) Slopegraphs summarizing the computing times of different recombination detection programs on alignments of 50, 100 and 200 sequences, derived from *n* = 37 HIV-1 subtype reference sequences with random mutations. We evaluated 10 replicate simulations per sample size. DSBM-8 and -12 correspond to dynamic stochastic block model analyses with 8 and 12 clusters, respectively. Note that DSBM computing times do not include the fast calculation of Tamura-Nei (TN93) distances. GARD was run in an MPI environment with 8 cores.

In addition, we evaluated the sensitivity of DSBM to varying the sizes of windows (250, 500, 750, 1000nt) and steps (50, 100, 200, 300nt), which determines the number and stability of the graphs generated from the genome alignment. Error percentage was not significantly associated with window size (quasibinomial GLM, *t* = − 0.33, *P* = 0.74) or step size (*t* = 0.75, *P* = 0.46), although we noted a slight tendency for error to increase with window size for simulations with 1 breakpoint, and to decrease with window size for 3 breakpoints (Supplementary Figure S2).

Finally, we measured the runtimes for processing varying numbers of sequences derived from HIV-1 subtype reference genomes with DSBM, GARD, RDP4 and RDP5 (Figure 1B). Since we were more interested in computing time than accurately inferring unknown recombination events in this experiment, we arbitrarily set the number of DSBM clusters to 8 and 12. Computing times for DSBM were comparable to RDP4, whereas GARD and RDP5 were substantially faster; however, it is difficult to compare the latter two since GARD requires a parallel computing (MPI) environment. Overall, these results indicate that the expected computing times tend to increase faster than linearly with the number of sequences, and increase with the number of clusters for DSBM. We also noted an unusual trend in GARD computing times, with a faster times obtained for sample sizes above 50 sequences. On examining the output files, we determined that GARD efficiently rejected all models with recombination breakpoints for the larger datasets. In contrast, GARD runs on datasets with 50 sequences were unable to reject the presence of recombination and were burdened with 677.8 candidate models (partitions of the alignment among trees) on average.

### Application to HIV-1/M genomes

Having validated the DSBM method on simulated data in comparison to other unsupervised recombination detection methods, we next applied our method to *n* = 525 full-length HIV-1/M genome sequences to characterize the role of recombination in the evolutionary history of this virus (Figure 2). These sequences were selected from all available full-length HIV-1 genomes to maximize the representation of global diversity in a reduced data set. The integrated completed likelhood (ICL) criterion, used to determine the optimal number of clusters for DSBM, was maximized at 25 clusters (Supplementary Figure S4).

**Figure 2:**
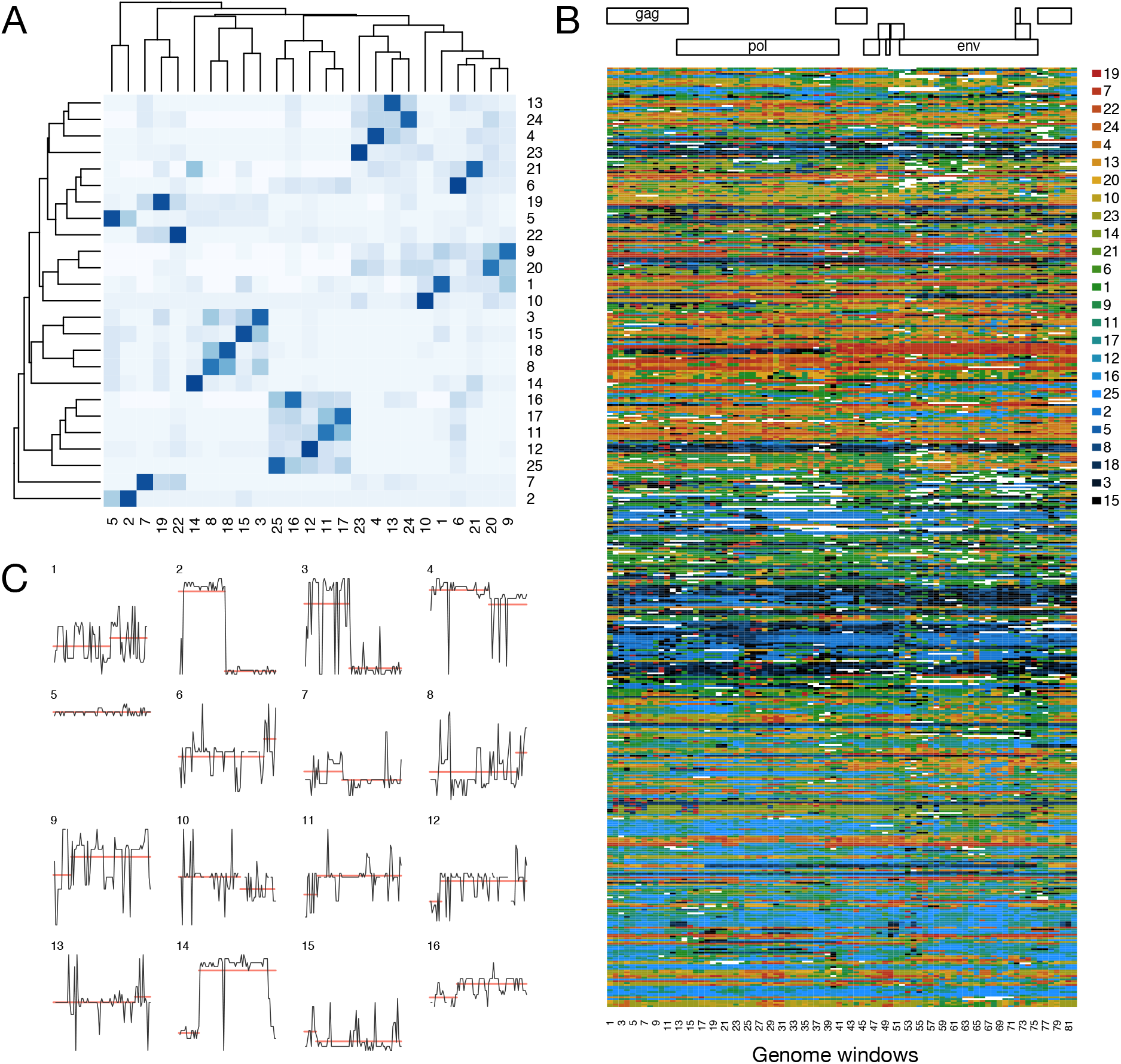
Summary of results from DSBM analysis of *n* = 525 HIV-1/M genomes. (A) A heatmap of the transition rate matrix estimated from the sequence of networks generated by a sliding window analysis of the HIV-1/M genome alignment. The highest rates (darkest shaded cells) corresponded to the rate that an individual remains in the same network community (cluster) on moving from one genomic window to the next. (B) A heatmap displaying the cluster assignments for *n* = 525 genomes, partitioned into *n* = 82 windows of 500nt in steps of 100nt. The colour map corresponds to the stacked barplot legend in (B). White cells indicate that DSBM failed to assign the respective genome to a cluster in that window. Distinct transitions between colours along a row implies recombination. A color-blind accessible version is provided as Supplementary Figure S3. (C) Stepcharts depicting the cluster assignments for the first 16 genomes across 82 windows, where we used the hierarchical clustering permutation order from (A) to minimize the vertical distance between clusters related by higher transition rates. Line segments (red) indicate the location of change points, as determined by a conservative ‘at most one change’ (AMOC) method.

The results of our analysis are summarized in Figure 2. Figure 2A displays a heatmap summarizing the transition rate matrix from the DSBM analysis. This rate matrix was not constrained to be symmetric, since the outflow from cluster 1 to 2 is not necessary balanced by a return flow from cluster 2. Some clusters have substantial transition rates (darker shades) between them, such as clusters 8 and 18, 11 and 17, or 16 and 25. This implies a hierarchical structure to clusters akin to the sub-subtypes of HIV-1. The frequencies of cluster memberships were consistent across the length of the HIV-1 genome (Supplementary Figure S5). To better understand the associations between these clusters, we mapped the original HIV-1 subtype annotations of the genome sequences to these data. We found that windows from genomes labelled as subtype A1 (*n* = 33) were predominantly assigned to clusters 4, 13 and 24; subtype B (*n* = 56) was associated with clusters 3, 8, 15 and 18; and subtype C (*n* = 66) to clusters 11, 16, 17 and 25 (Supplementary Figure S6). Within these major subtypes, cluster assignments tended to become more variable past genome window 50 (Supplementary Figure S7), in association with the *env* gene. In addition, subtypes D, G, F1 and the circulating recombinant form (CRF) 01AE associated strongly with clusters 14, 10, 2 and 7, respectively. Clusters 1 and 9 associated with subtypes J, H and K, as well as genomes that were annotated as ‘complex’ inter-subtype recombinants.

Figure 2B displays the cluster memberships for all 525 genomes in 82 windows of 500nt in steps of 100nt. We mapped cluster memberships to a colour gradient such that clusters with similar colours tend to have relatively higher transition rates between them. Under the standard concept of ‘pure’ HIV-1 subtypes, genomes should tend to maintain the same colouration across its length. Indeed, we see an overall stratification of colours in this heatmap (Figure 2B). However, if we examine the cluster assignments for individual genomes (Figure 2C), we see traces that are consistent with recombination, such as genomes 2, 3 and 14. Other traces are too noisy to be interpreted visually. Therefore, we used two different change point detection algorithms to quantify the extent of recombination from the entire set of traces. The AMOC method classified 465 (95.9%) genomes as recombinant (at least one change point) and 20 as non-recombinant (40 genomes were excluded due to excessive missing values). This result was robust to varying the minimum segment length; for example, AMOC only classified 47 (9.7%) genomes as non-recombinant at a cutoff of 20 windows.

Next, we used the less restrictive PELT method to estimate the number of breakpoints per genome (Figure 3, left). As expected, the mean number of breakpoints per genome increased with shorter minimum segment lengths. At a minimum length of 5 windows, for instance, we inferred a mean of 6.8 (IQR 5-12) breakpoints. On the other hand, the distribution of breakpoints across windows was less sensitive to varying the minimum segment length, with significant positive correlations between distributions (Spearman’s *ρ >* 0.58, *P* < 4.7 *×* 10^− 7^; Figure 3, right). Local peaks in the frequencies of breakpoints were robust to varying minimum segment length. Consistent with previous work [50], we observed recombination ‘hotspots’ associated with the 5’ and 3’ ends of the *env* gene. We also observed a distinct and robust peak associated with the 3’ end of the *pol* gene, and a sharp increase in the frequency of breakpoints within or upstream of *gag*. Genomes that were classified as recombinant by SCUEAL had significantly higher numbers of breakpoints assigned by DSBM, and this concordance was robust to varying minimum segment lengths (Wilcoxon rank-sum test, *P* < 6.3 *×* 10^−3^; Supplementary Figure S9). Furthermore, we found no significant association between the number of predicted breakpoints and the year of sample collection, which ranged from 1983 to 2016 with a median of 2006 (linear model *t* = 0.86, *P* = 0.39; Supplementary Figure S8).

**Figure 3:**
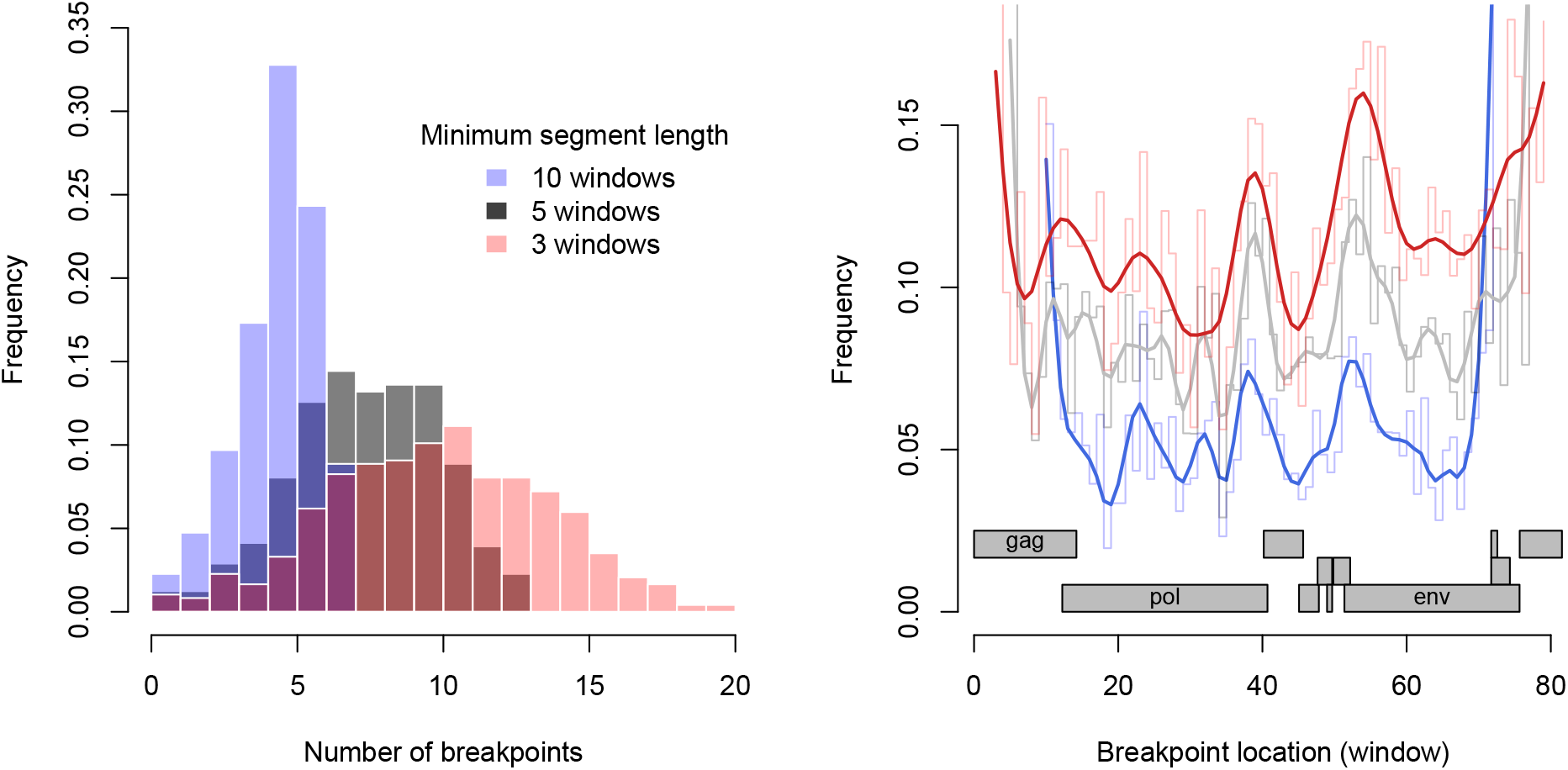
Distribution of DSBM predicted recombination breakpoints in HIV-1/M genomes. (left) Histogram of the number of breakpoints per genome, where breakpoints were extracted from DSBM cluster assignments by the PELT change detection method. We varied the minimum segment length to 3, 5 and 10 windows (see colour legend). (right) Smoothed splines summarizing the distribution of predicted breakpoints among sliding windows of the HIV-1 genome alignment. We used the same colours as the histograms to indicate minimum segment lengths. The raw frequency distributions are drawn as step charts in lighter colours. A diagram of the HIV-1 reading frames, mapped from the HXB2 reference coordinates to our alignment, is displayed at the base of the figure.

### Revisiting the HIV-1 subtype references

In addition to analyzing a large selection of HIV-1/M genome sequences, we applied the DSBM method to evaluate the HIV-1 subtype reference genomes curated by the Los Alamos National Laboratory HIV Sequence Database. These reference genomes have long been used as the ‘gold standard’ against which other genome sequences are evaluated for evidence of recombination. We assessed a subset of the reference genomes covering HIV-1/M subtypes A-D, F-H, J and K, and excluded reference genomes corresponding to the circulating recombinant forms (CRFs). For this analysis, the ICL criterion selected *k* = 6 as the optimal number of clusters. This is clearly fewer than the recognized number of HIV-1 subtypes. We observed that subtypes B and D were assigned to the same cluster. Furthermore, genomic windows from subtypes F and K tended to be assigned to the same cluster, and subtypes G and J to a third cluster. These cluster assignments are consistent with the placement of the respective subtypes in a phylogeny of HIV-1/M subtypes and CRFs [51].

Employing the same post-processing method as our previous analysis, we detected evidence of recombination breakpoints in 18 (48.6%) of 37 reference genomes when the minimum segment length was set to 5 windows. The PELT method of change point detection identified 1, 2 and 4 breakpoints in 2, 13 and 3 of the remaining genomes, respectively (Figure 4). These breakpoints tended to be concentrated in genomes classified by SCUEAL into subtypes F (sub-subtypes F1 and F2), J and K. We also observed consistent patterns of recombination in the subtype G genomes spanning windows 42-51 and 67-78. These putative breakpoints, which are consistent with previous work [19], were not picked up by the PELT method because the affected clusters were annotated as being similar, even though we did not observe substantial clustering in transition rates between the 6 clusters in this analysis. These results highlight an important limitation of change point detection based on changes in the mean.

**Figure 4:**
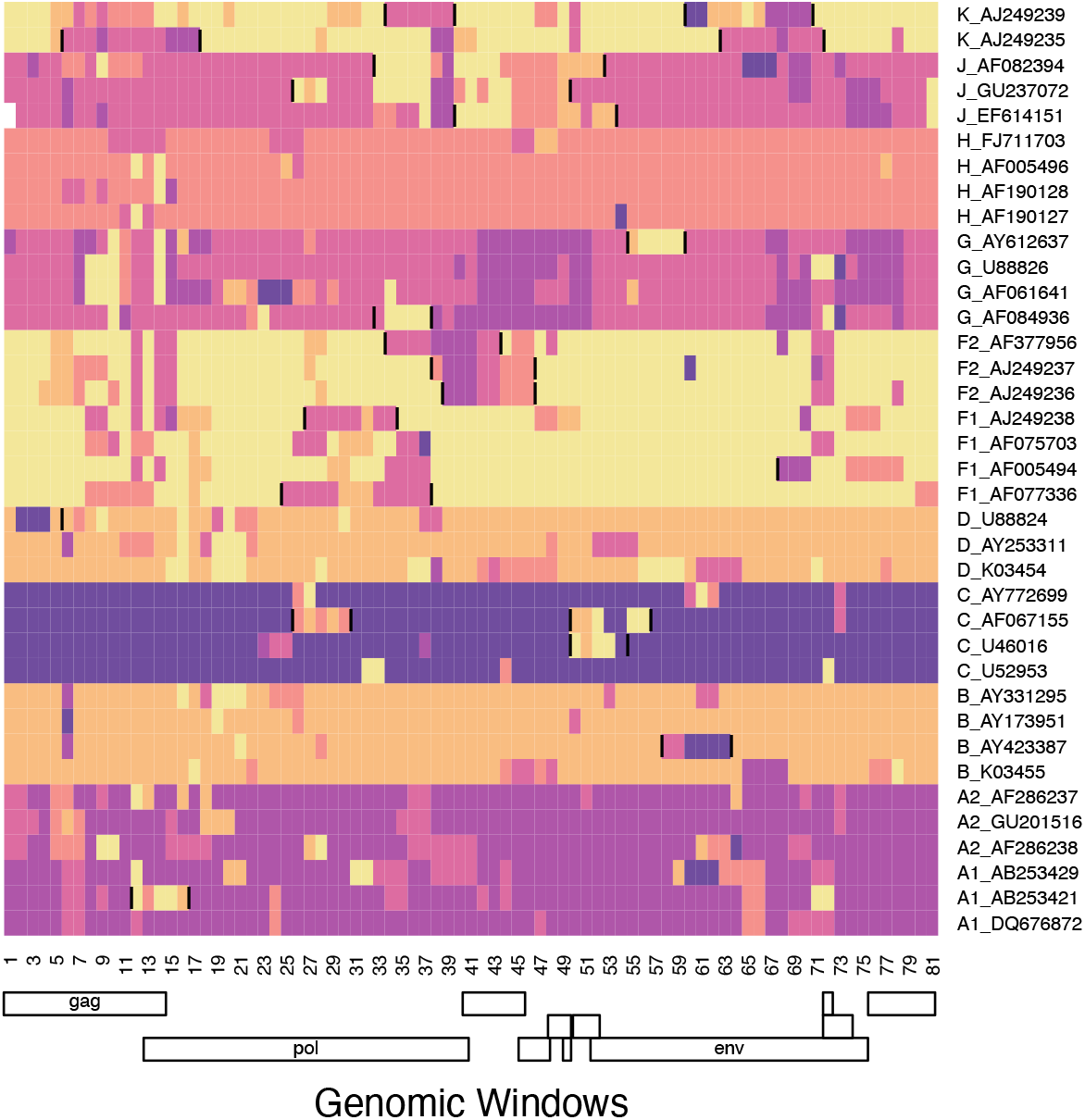
Detection of recombination in curated HIV-1 subtype reference genomes by community detection with a dynamic stochastic block model (DSBM). Predicted breakpoints (using the PELT change point detection method with a minimum segment length of 5 windows) are marked directly on the heatmap with vertical line segments. We used a different colour palette to emphasize the different number of clusters between this analysis (*k* = 6) and our previous analysis of *n* = 525 HIV-1/M genomes (*k* = 25).

## Discussion

Recombination is a major contributing factor to the extensive genetic variability observed in RNA viruses [52]. Therefore, it is pertinent to screen for recombination when studying evolution in these viruses [22]. Recent advancements in whole genome sequencing is driving the proliferation of large HIV-1/M genome databases [53, 54], providing more opportunities to reconstruct the role of recombination in the evolutionary history of this virus. Here, we have adapted a community detection method from network science (dynamic stochastic block models, DSBM [24, 32]) to detect the residual evidence of past recombination events without requiring a reference set of non-recombinant genomes. Hence, we characterize DSBM as an unsupervised method for detecting recombination.

Compared to the commonly-used unsupervised recombination detection programs GARD, RDP4 [49] and RDP5 [23]. GARD employs a genetic algorithm to explore the model space of placing one or more recombination breakpoints in the alignment, reconstructing a maximum likelihood phylogeny for each interval between breakpoints [22]. Breakpoints are accepted if the increase in likelihood is sufficient to justify increasing the number of model parameters induced by the addition of a discordant phylogeny. Of the unsupervised methods, it is the closest to a parametric model of the evolutionary consequences of recombination. RDP versions 4 and 5, on the other hand, employ a suite of non-parametric recombination detection methods. For example, BOOTSCAN calculates pairwise distances between sequences in sliding windows. Thus, our initial step of mapping a sequence alignment to DSBM is similar to RDP. On the other hand, DSBM operates on a more comprehensive representation of sequence diversity by converting the pairwise distance matrices into a series of undirected graphs. In addition, DSBM employs a hidden Markov model in which the probability of an edge connecting two individuals is determined by their unobservable (latent) and dynamic cluster memberships. Consequently, the placement of a breakpoint at a particular location in one sequence is informed by observed patterns across all sequences.

DSBM was significantly more accurate for detecting recombination in simulated data. However, DSBM is time-consuming to compute and slower than the other methods, in part due to the quadratic time complexity of parameterizing the matrix of transition rates between clusters (communities). Like GARD, the computing time can be ameliorated by running DSBM in a parallel computing environment. Our extension of DSBM involves a number of tuning parameters — namely, a genetic distance threshold, the window size, and the window step size. Our simulation experiments indicated that results from DSBM are not sensitive to varying the window and step size parameters. However, choosing an appropriate distance threshold, which determines how sequence variation maps to network topologies, was critical to our analysis (Supplementary Figure S1). Another limitation of DSBM is that it assumes the rates of transitions between clusters are ‘time’ homogeneous, *i*.*e*., that recombination rates between specific lineages are constant through the genome. Since DSBMs are a recent innovation in network science, it may eventually become possible to fit models with time-heterogeneous rates; however, the current model already estimates a large number of parameters from the data.

An interesting outcome of our analysis is the putative recombination hotspots in association with the *gag* and *nef* gene sequences, which are adjacent to or overlap the 5’ and 3’ long terminal repeats (LTR), respectively (Figure 3). Based on the composition of the CRF genomes documented by the Los Alamos National Laboratory HIV Sequence Database, breakpoints are often found in these regions. For example, CRF01 AE contains a breakpoint at the 5’ end of the 3’ LTR sequence (HXB2 nucleotide coordinate 9086), and CRF02 AG at the 3’ end of the 5’ LTR (coordinate 789). Overall, we observed that 57.1% (*n* = 77) and 45.6% (*n* = 92) of the documented CRFs contain breakpoints upstream of HXB2 coordinates 1496 and 8757, respectively (adjusting for CRFs without sequence coverage in these regions). Furthermore, recombination in these regions of the HIV-1 genome has been observed as non-CRFs [55, 56]. In a comparative analysis of HIV-1 *gag*, Minin *et al*. [57] detected a recombination hotspot in association with an instability element in the region encoding the capsid protein. However, this hotspot was not reproduced in subsequent work by Archer *et al*. [50], who reported hotspots associated with both ends of the *env* gene. Resolving these differences will require the reconciliation of methods and data sets, or more effectively, experimental validation with an *in vitro* system [58].

Previous studies have proposed re-classifying some of the defined HIV-1 subtypes and circulating recombinant forms. In some cases, this was motivated by the availability of new genomic data [59, 60]. For example, a recent analysis of the major HIV-1 genes *gag, pol* and *env* culminated in a proposal to further partition of subtypes A and D into sub-subtypes, and to merge subtypes B and D into a single subtype [61]. Moreover, a new non-recombinant HIV-1/M subtype L was proposed almost 30 years after the genomes were first sampled in 1983 and 1990 in what is now the Democratic Republic of the Congo [62]. Our analysis indicates that many of the HIV-1/M subtype reference genomes are actually recombinant, which is consistent with previous work [18, 19]. These findings support the idea that the current HIV-1 nomenclature [7], which has served as an important framework for our understanding of HIV-1 diversity and evolution, should be revisited in light of new genomic evidence. Our DSBM analysis of a large sample of HIV-1/M genomes identified *k* = 25 as the optimal number of clusters. In this context, a cluster is roughly analogous to a non-recombinant subtype. However, what we recognize as a subtype may also comprise multiple clusters (Supplementary Figure S7). None of the *n* = 525 in our largest analysis was assigned along its entire length to a single cluster. It is unlikely that expanding the scope of our analysis will yield genomes that are each representative of a single cluster. However, it may be feasible to utilize the cluster assignments from DSBM to generate composite ancestral genome reconstructions that can play the same role as the current HIV-1 subtype reference genomes.

## Acknowledgements

This work was supported in part by the Government of Canada through Genome Canada and the Ontario Genomics Institute (OGI-131) and by the Canadian Institutes of Health Research (CIHR grants PJT-155990, PJT-156178, FRN-130609, BOP-149562).

## Supplementary Figures

**Figure S1:**
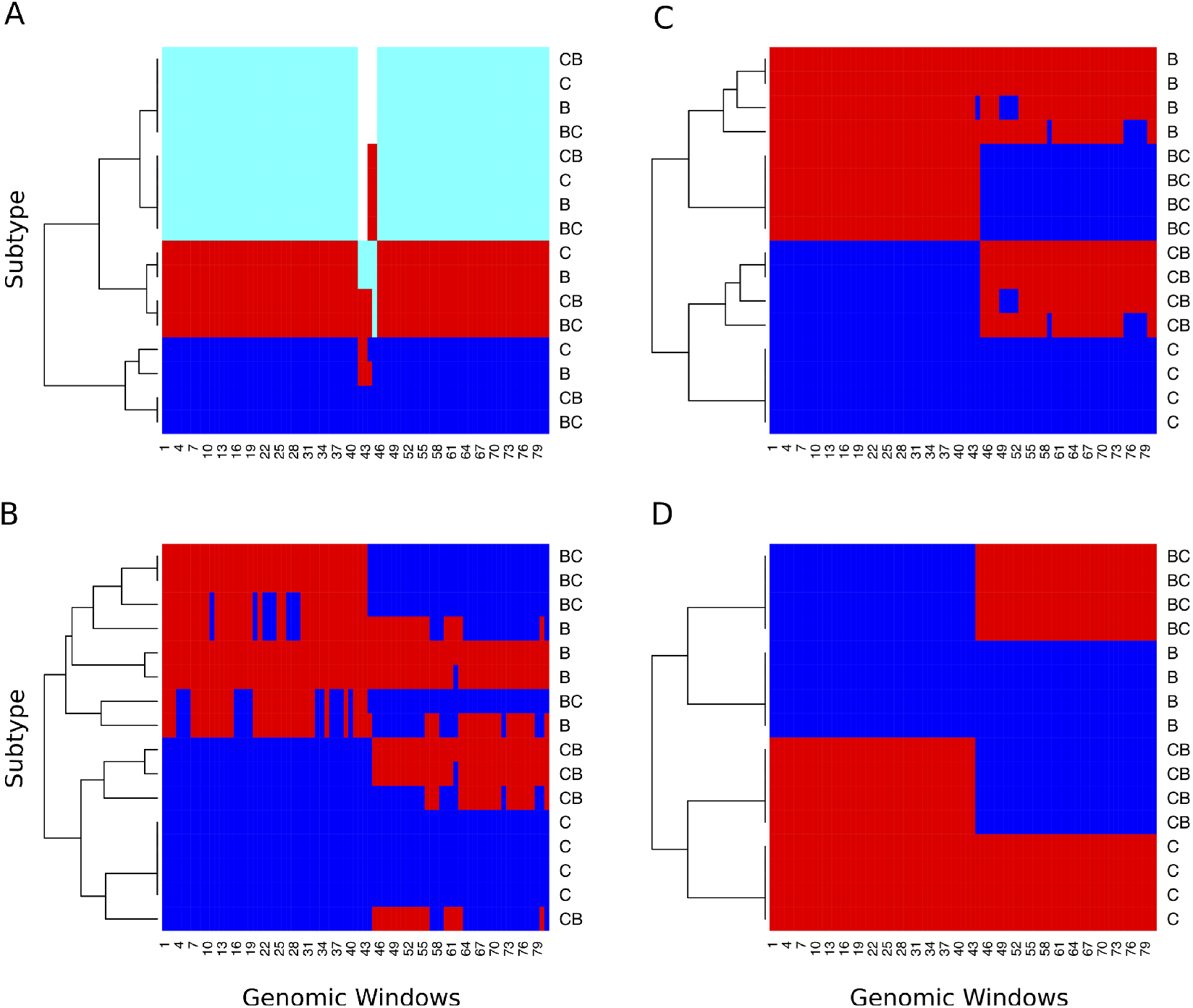
An example of DSBM cluster assignments on simulated HIV-1 recombinant genomes, displayed as heatmaps. We generated graphs from an alignment of *n* = 16 HIV-1 genomes for varying Tamura-Nei (TN93) distance cutoffs: A=10%, B=20%, C=30%, D=40%. Each row in the heatmap represents a genome sequence, and each column represents one of *n* = 82 sliding windows of 500nt. Each cell in a heatmap is coloured with respect to the DSBM cluster assignment, with the optimal number of clusters determined by the ICL criterion. The ‘true’ labels of the genomes are indicated on the right-hand side of each heatmap.

**Figure S2:**
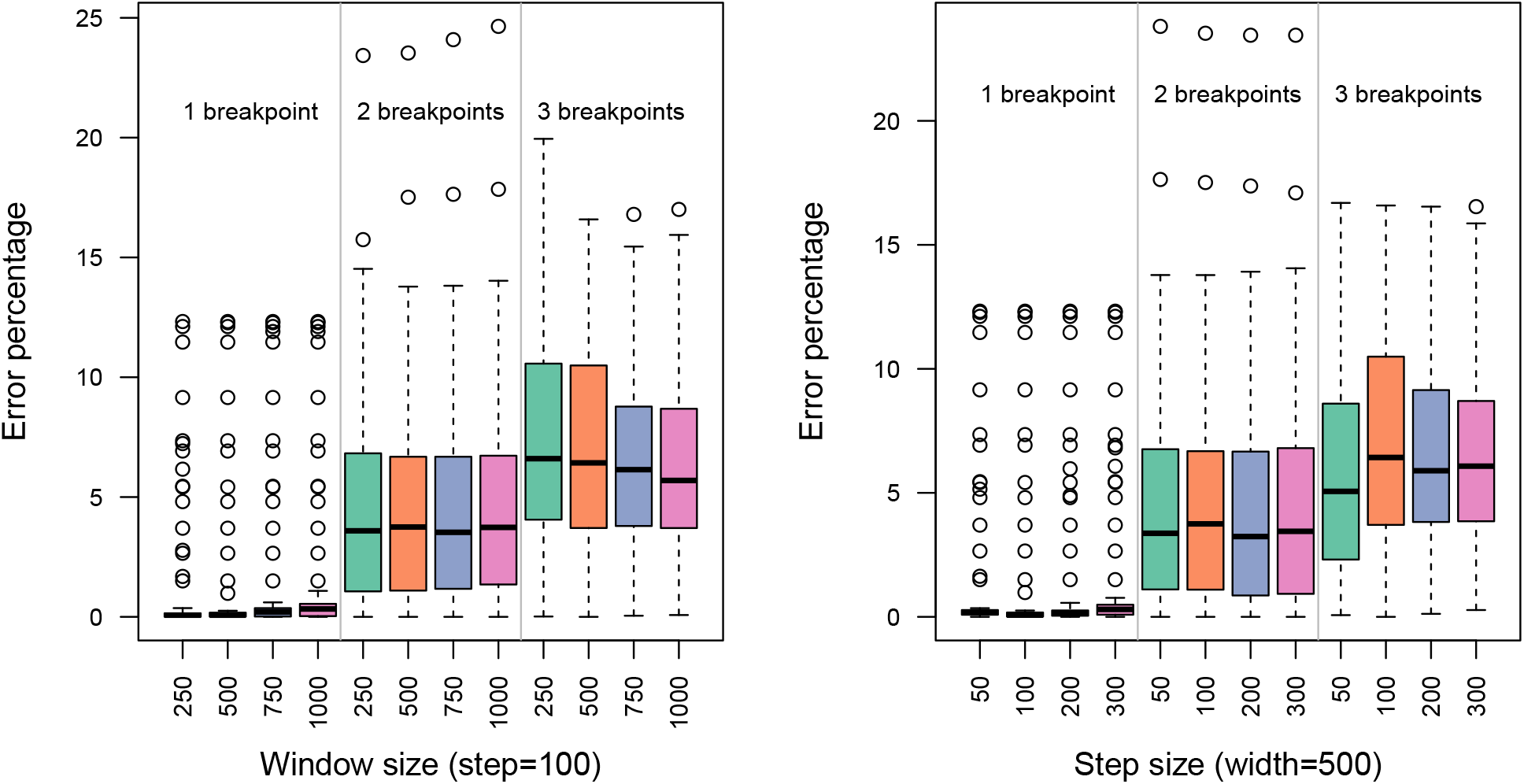
Box-and-whisker plots showing the percentage error reconstructing simulated recombination breakpoints using DSBM with varying window (left) and step (right) sizes.

**Figure S3:**
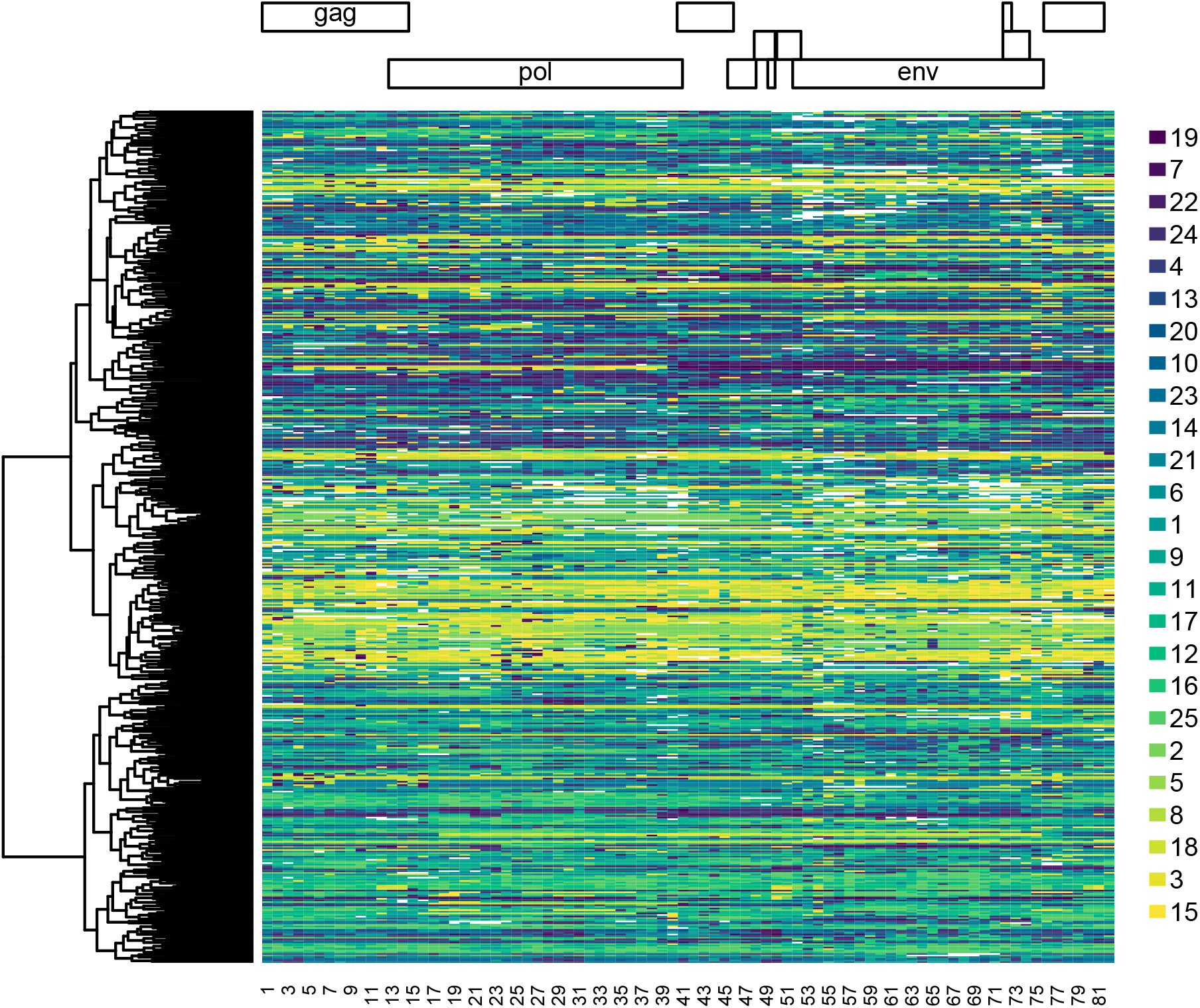
Color-blind accessible (viridis palette) version of heatmap summarizing DSBM cluster assignments for HIV-1/M genome sequences (Figure 2B).

**Figure S4:**
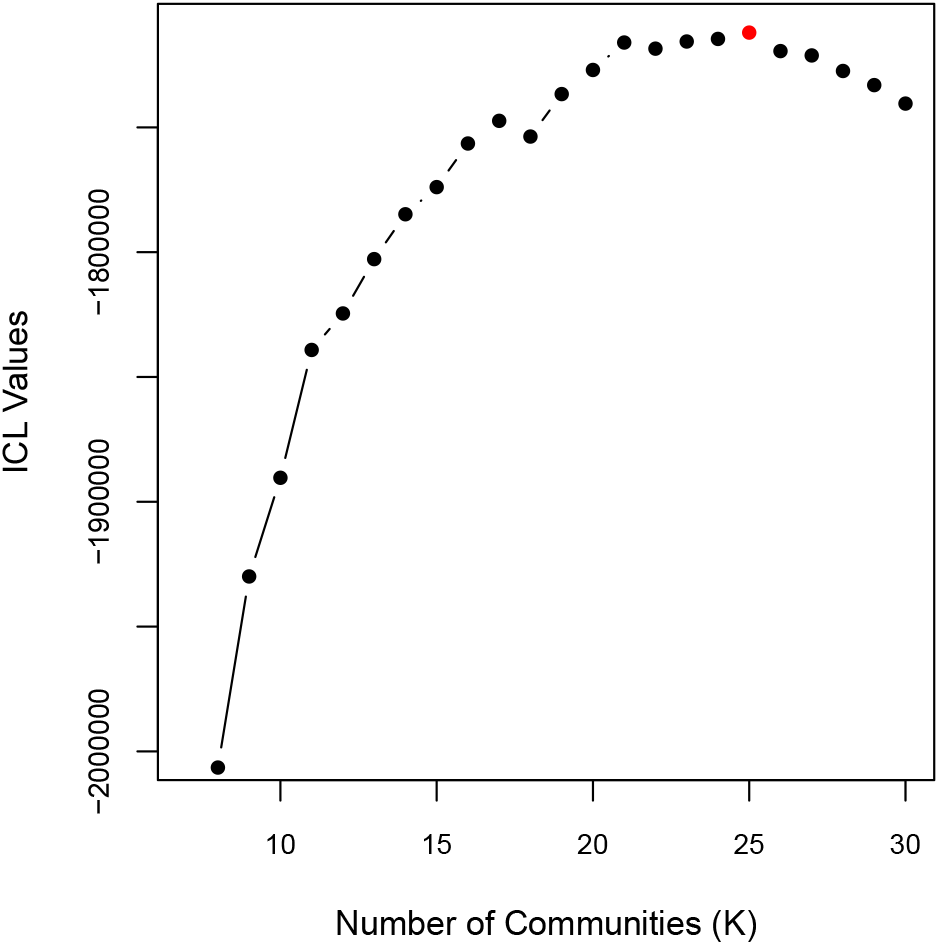
Profile of integrated composite likelihood (ICL) values as a function of *k*, the number of network communities, in DSBM analyses of graphs derived from the HIV-1/M genome windows. The maximal value (*k* = 25) is highlighted in red. Lower values of *k* were truncated from the plot to clarify differences in ICL in the neighbourhood of the maximal value.

**Figure S5:**
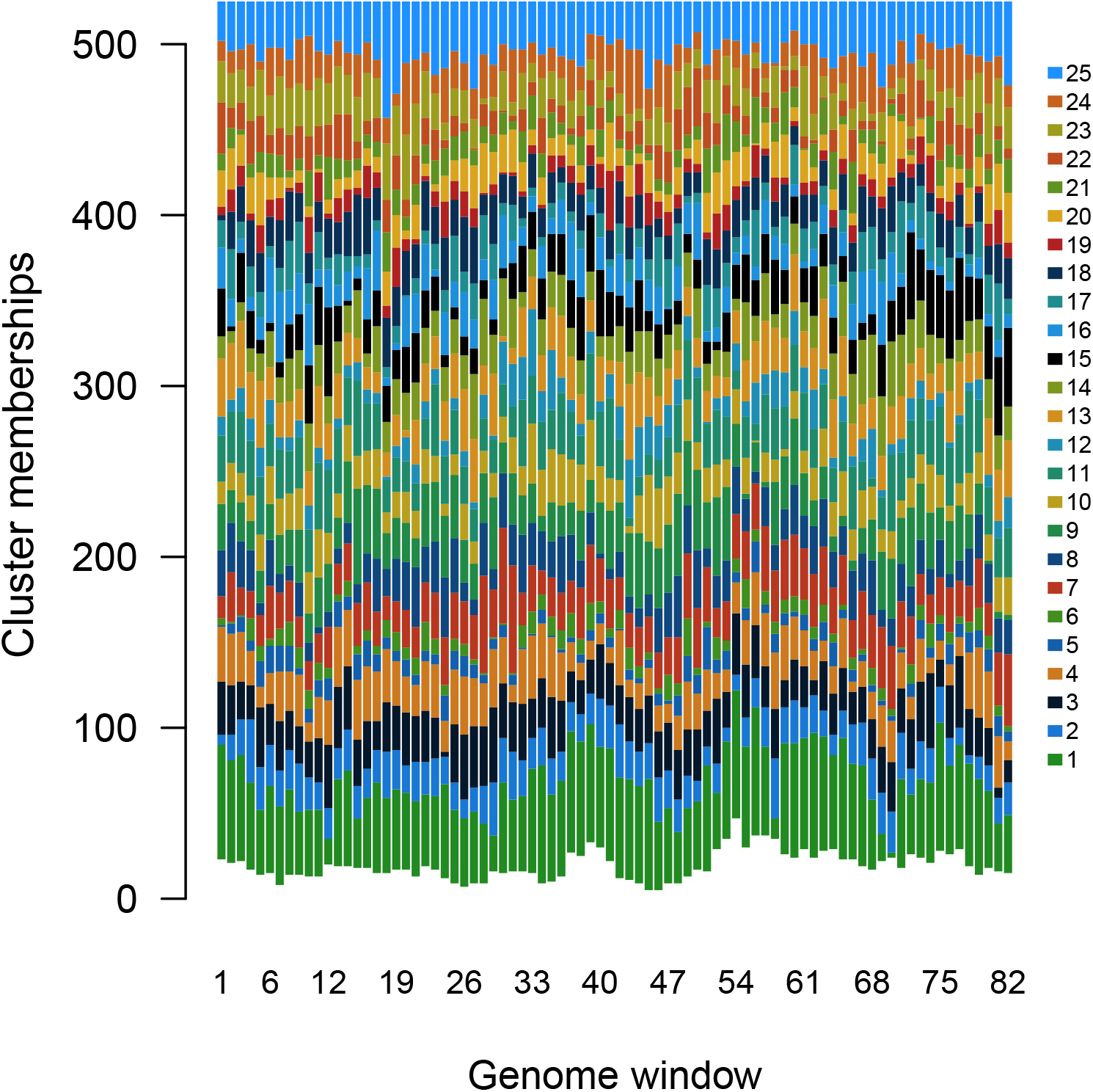
This stacked barplot summarizes the frequencies of cluster memberships across genome windows. We mapped a colour gradient to the *k* = 25 DSBM clusters using the permutation order from the hierarchical clustering used to generated the heatmap, such that clusters connected by higher transition rates are assigned similar colours. The bars have different heights because a varying number of genomes could not be assigned to any cluster (index 0) in a given window.

**Figure S6:**
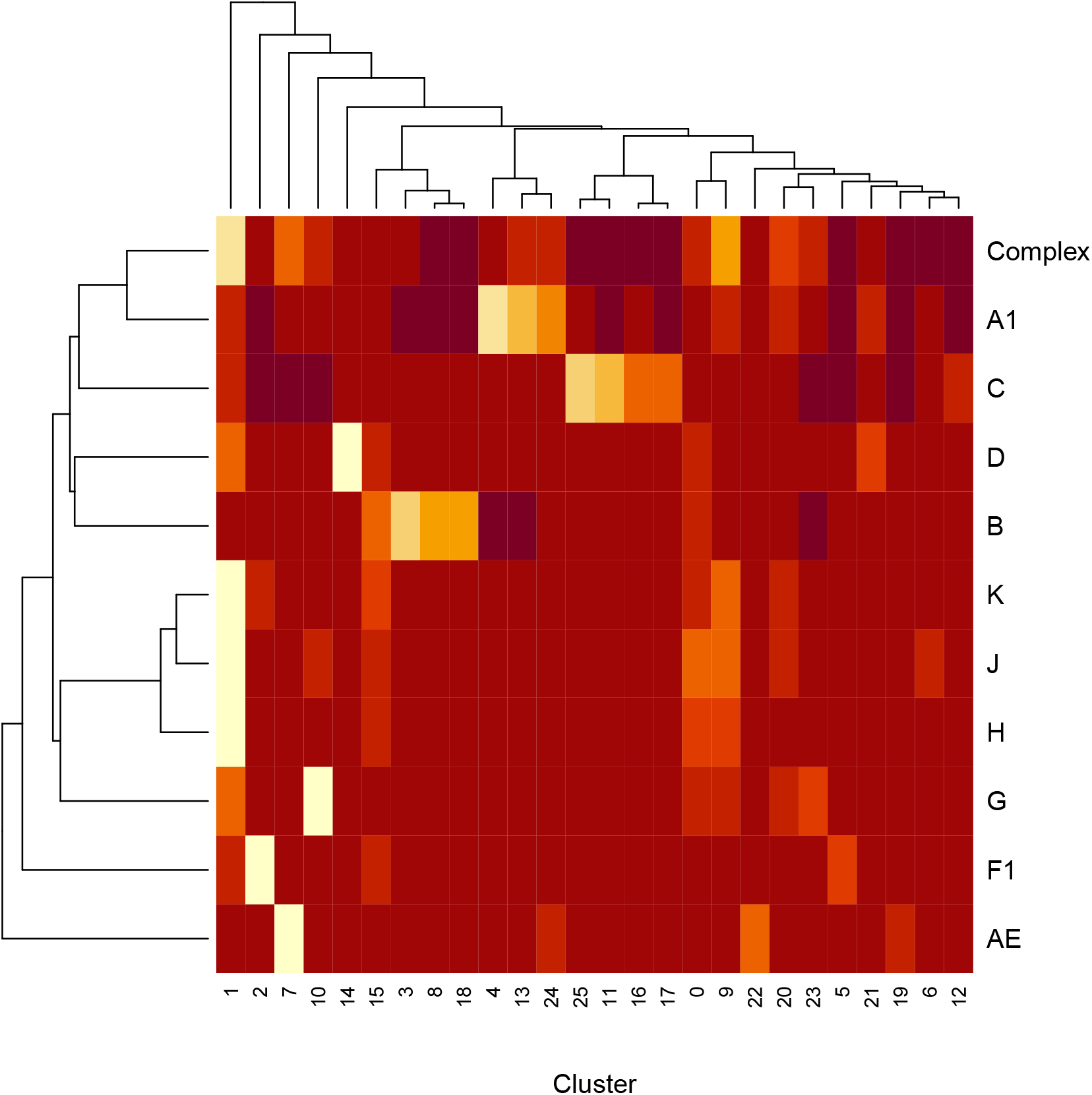
Heatmap summarizing the distribution of cluster assignments among HIV-1 subtypes. Subtype designations were generated using the SCUEAL (subtype classification using evolutionary algorithms) method as implemented in the software package *HyPhy*. White indicates higher frequencies of cluster assignments, whereas dark red indicates lower frequencies.

**Figure S7:**
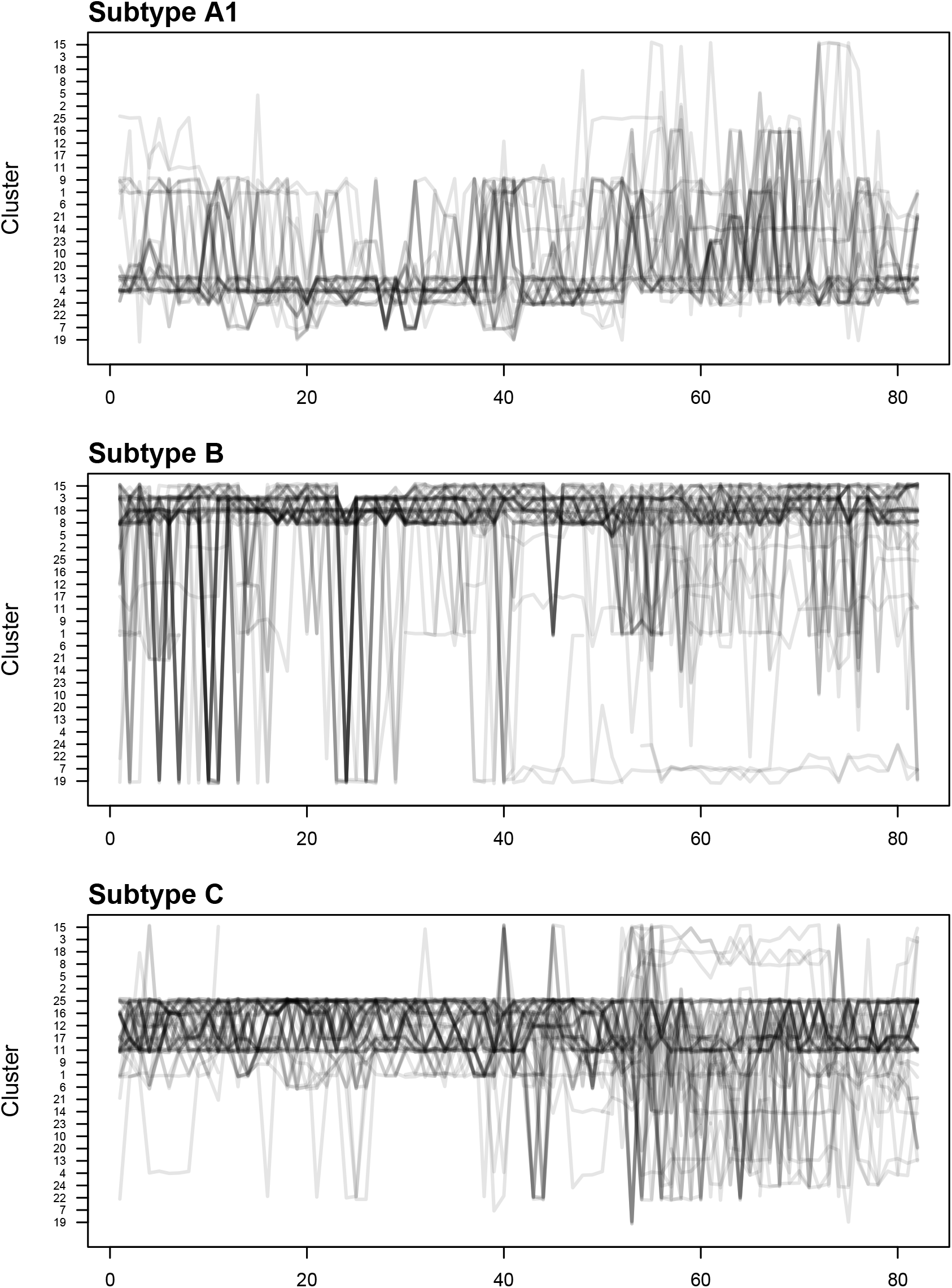
Plot showing community transitions in 3 pure subtypes (A1, B and C).

**Figure S8:**
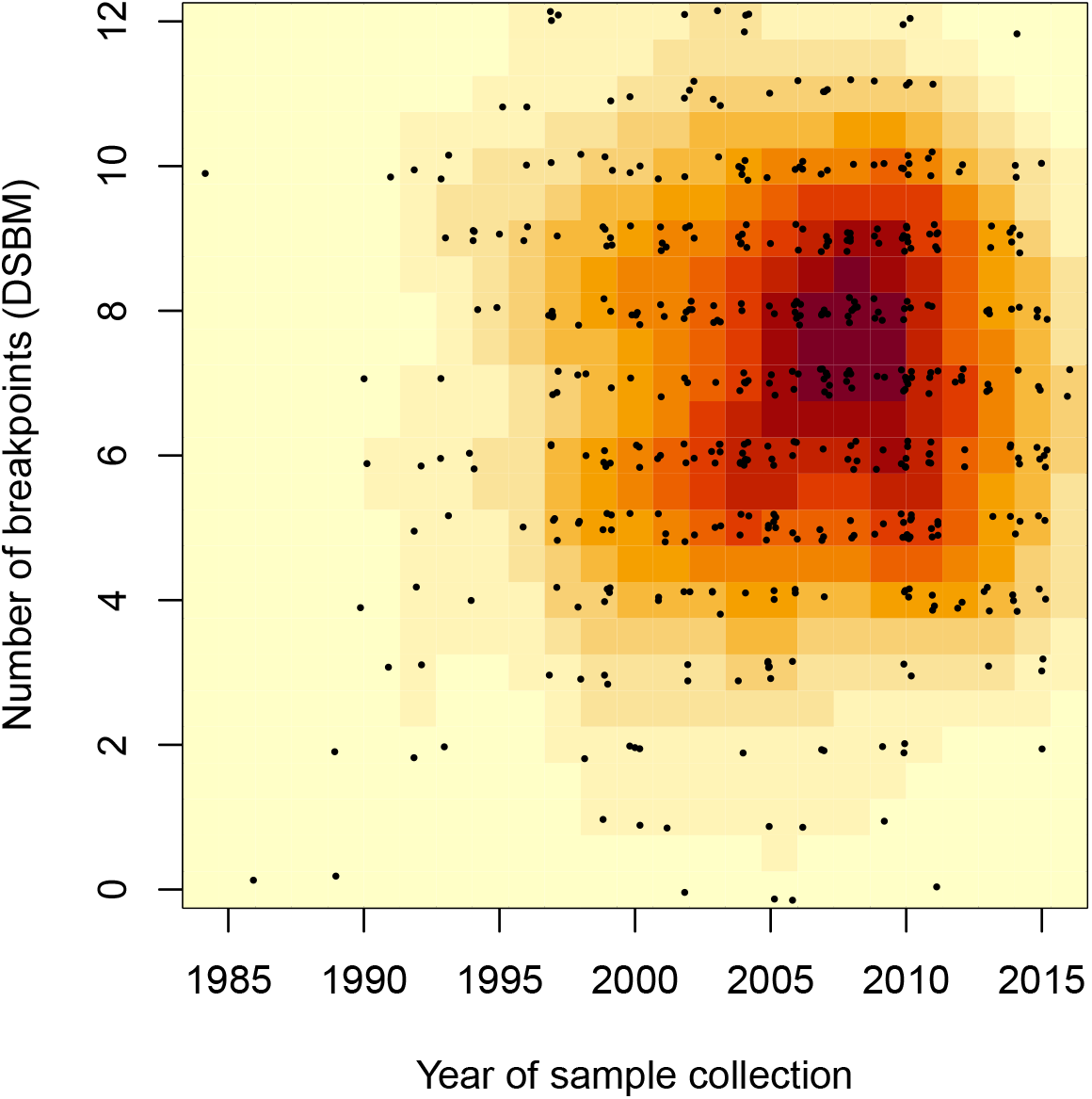
Distribution of DSBM-predicted recombination breakpoints in HIV-1/M genomes over time. This plot summarizes the distribution of genomes with respect to the number of breakpoints and year of sample collection with a kernel density, mapped to a heat colour palette with dark red for higher densities. In addition, the individual data points are mapped to the same plot region with added random noise with respect to both variables to reduce overlap.

**Figure S9:**
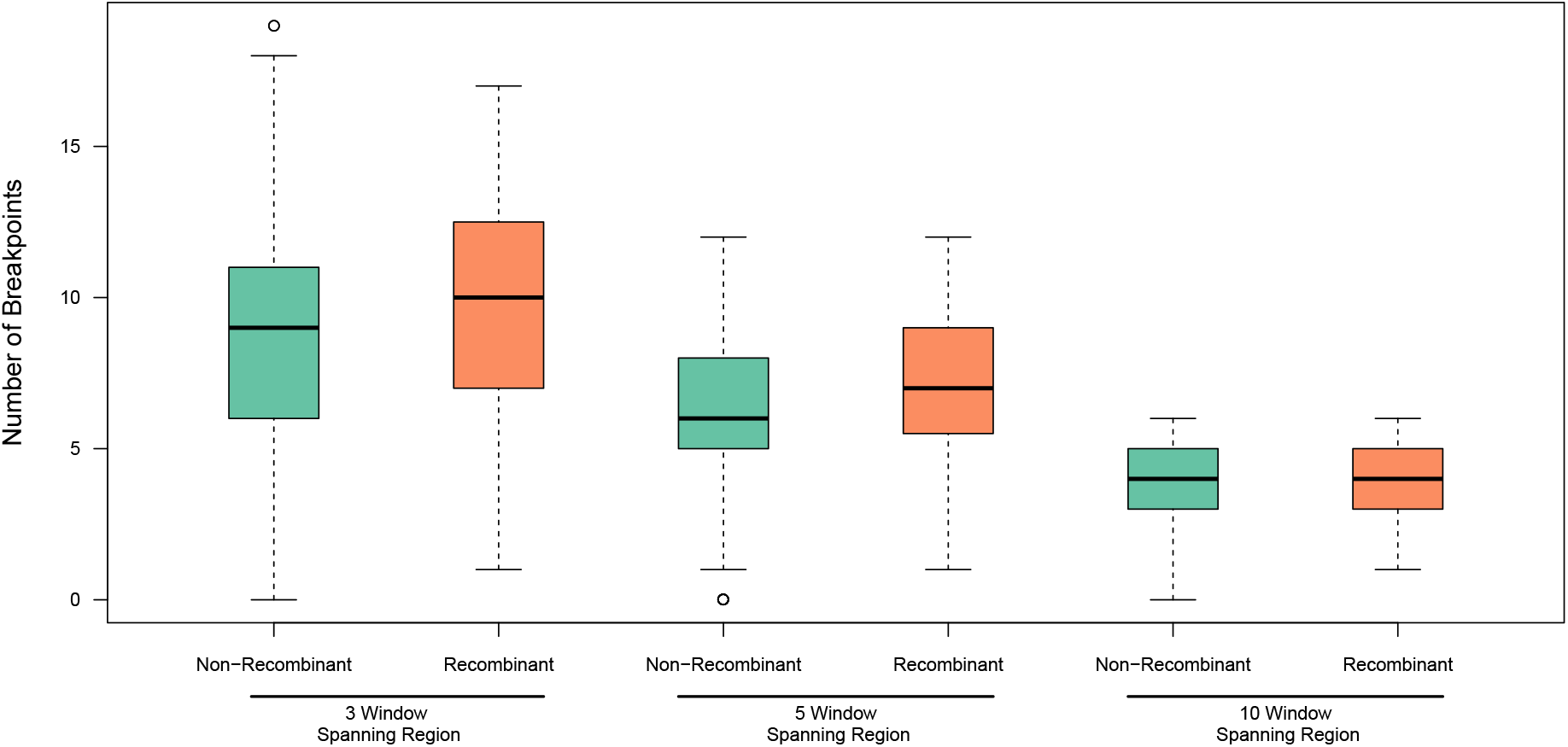
Frequency distribution of breakpoints between non-recombinant and recombinant genomes as classified by SCUEAL.

